# Food web structure mediate positive and negative effects of diversity on ecosystem functioning in a large floodplain river

**DOI:** 10.1101/2024.01.24.576859

**Authors:** Dalmiro Borzone Mas, Pablo A. Scarabotti, Patricio Alvarenga, Pablo A Vaschetto, Matias Arim

## Abstract

Research programs on Biodiversity-Ecosystem Functioning (BEF) and Food Web Structure (FWS) have contributed to understanding the impact of biodiversity on the functioning and architecture of ecosystems, but the interconnectedness between these components was seldom attended until recently. Several theoretical hypotheses predict an interconnection between BEF and FWS but were poorly and independently evaluated. We estimated 63 sink food webs of predatory fish in the Paraná River, covering a large gradient of community richness. We evaluated available hypotheses and their interrelationship through path analyses. A well-supported causal structure was identified, supporting that species richness directly increased standing biomass, modularity, and intermodular connection, whereas decreased interaction strength, connectance, and nestedness. A direct positive effect of modularity and connectance on biomass indicates that FWS can determine the BEF. Richness promotes biomass directly and through the increase in modularity but can also decrease biomass due to the decay in connectance, with similar positive and negative effects of richness on biomass. In this sense, the relationship between diversity and ecosystem functioning cannot be blind to FWS. Environmental homogenization and reduction in functional diversity may undermine the conditions for modular food webs, switching positive BEF to negative ones with potential cascading effects in the whole ecosystem.

## Introduction

The current biodiversity crisis has raised concerns about the impacts of the decline in species richness on the provisioning of ecosystem services, making the relationship between diversity and ecosystem functioning a key area of inquiry in ecology (Isbell et al. 2011, Tilman et al. 2014, Loreau et al. 2021). Many factors, including environmental conditions (Baert et al. 2018, Li et al. 2020, Ashford et al. 2021), functional diversity (Duffy et al. 2017, Maureaud et al. 2019), spatial processes (Thompson et al. 2018, Borthagaray et al. 2020), eco-evolutionary dynamics (Loeuille 2019, Akesson et al. 2021), and trophic interactions (Loreau & Hector 2001, Mora et al. 2014, Wu et al. 2022), have been identified as determinants of the relationship between species richness and ecosystem functioning (van der Plas 2019). However, in spite that a tight association between diversity and complexity has been largely recognized (Valdovinos et al. 2010; McCann 2011; May 1972; Thébault et al. 2010), the role of food web structure in the diversity - ecosystem function relationship has been largely overlooked (Nie et al. 2023). Indeed, recent research indicated that resource distribution due to food web interactions, cascading effects between predators and their prey, and different facets of food web complexity such as interaction strength or connectance, may shape the diversity-functioning relationship, supporting that this relationship cannot be studied in isolation from the structure of food webs (Carroll et al. 2011; Poisot et al. 2012, Mora et al. 2014, Yang et al. 2021, Wang et al. 2022; Nie et al. 2023). This framework demands for the advancement in the relationships between: 1) diversity-food web structure, 2) food web structure-ecosystem functioning, and 3) diversity - ecosystem functioning. Furthermore, it is crucial to consider all these components simultaneously to understand the direct and indirect mechanisms that shape the functioning of ecosystems (Loreau et al. 2010, Poisot et al. 2012; Wang and Brose 2018; Wang et al. 2022; Nie et al. 2023).

The relationship between diversity and food web structure raised much interest after the provocative studies of Robert May (1972) pointing that diversity by itself undermines stability, paving the way for a fruitful research program that is maturing in recent years (Valdovinos et al. 2010; McCann 2011). This program has established several well-known relationships, including the decrease in food web connectance with species richness (May 1972; Thébault et al. 2010), the increase in weak interactions with the ratio between predator-prey body size and prey diversity (Brose et al. 2006; Loeuille 2010; Brose et al. 2019, DeLong et al. 2021), the prevalence of modularity and its role in stabilizing ecological networks (Stouffer & Bascompte 2011), the progressive coupling of energy channels by predators in higher trophic positions (Rooney & McCann 2008; Arim et al. 2010), the network rewiring by adaptive foragers (Valdovinos 2010; Loeuille 2010; Bartley et al. 2019) the spatial arrangement of food webs in metacommunities (Pillai et al. 2012: Mougi and Kondoh 2016), and the decrease in resource use of generalist over specialist species (i.e. nestedness) with the increase of diversity (Trojelsgaard & Olesen et al. 2013, Pinheiro et al. 2019, Wen et al. 2022). All these structural properties are expected to become more important as more species are interacting in the community (Fontaine 2013, Welti & Joern 2015), promoting well-defined changes in food web structure as diversity increases (Fig. 1). Despite its central importance, a comprehensive empirical analysis of the entangled relationships among these metrics along a species richness gradient has been considered only in a few papers (e.g. Thébault et al. 2010; Nuwaga et al. 2017; Danet et al. 2021, Nie et al. 2023). The untangling of these relationships is a central challenge, because many structural properties of food webs are tightly interrelated and several alternative causal structures are congruent with the proposed association between diversity and food web metrics. For example, the association between metrics as nestedness-modularity (Fortuna et al. 2010), modularity-connectance (May 1974), interaction strength-nestedness (Basotlla et al. 2009), cast doubts about the metrics that are effectively responding to variations in richness. In addition, a direct effect of diversity in metrics as modularity or connectance will affect all the other metrics, highlighting the potential role of indirect effects (Eschenbrenner and Thébault 2023; Nie et al. 2023).

**Figure 1:**
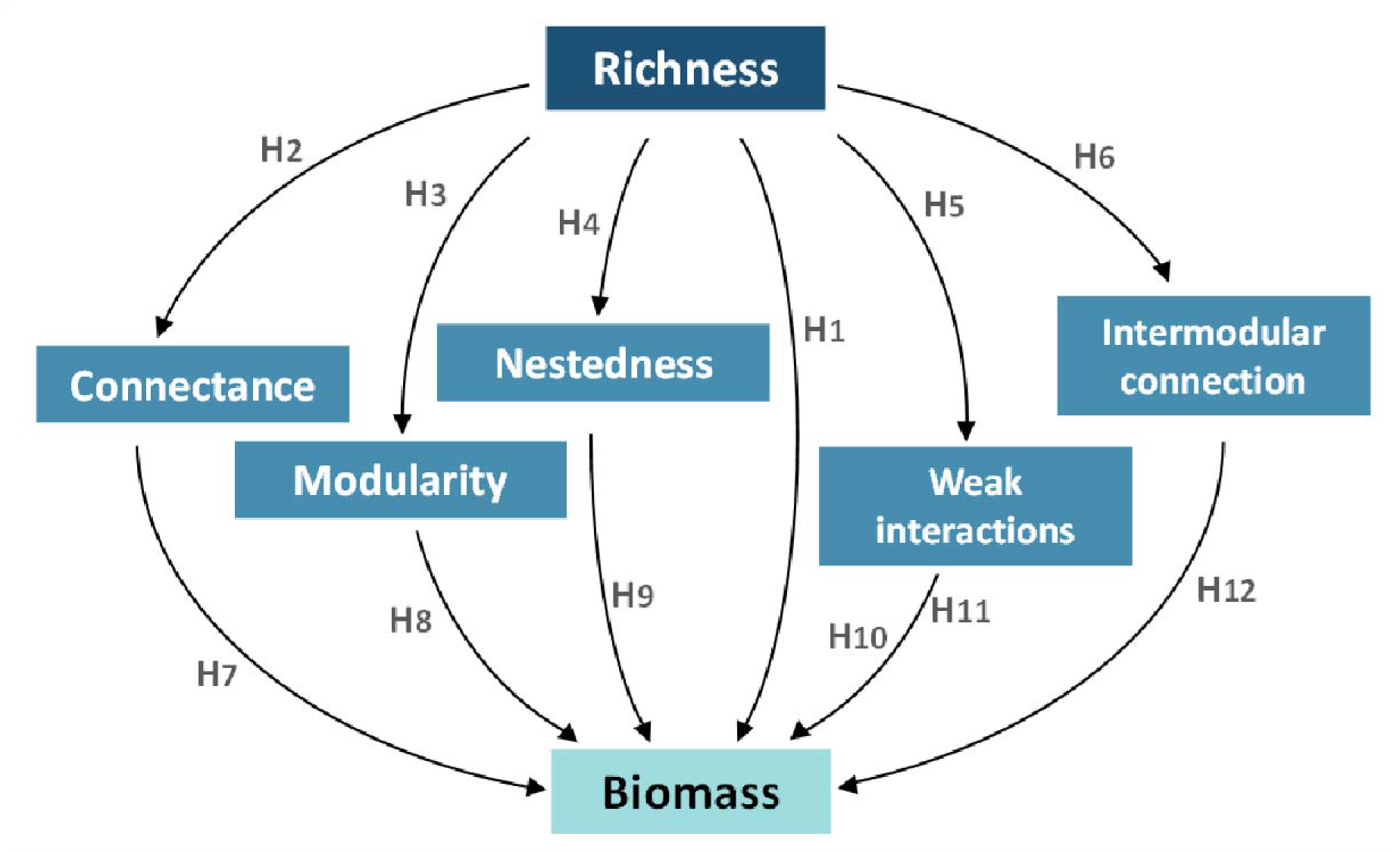
Hypothesized causal connection between community richness, food web structure and biomass production. The box text indicates the hypothesis or empirical evidence supporting each one of the links considered.

The relationship between diversity and ecosystem functioning has been a major focus of research in ecosystem theory (Loreau 2010; Tilman et al. 2014, van der Plas 2019), with numerous studies demonstrating a positive correlation between diversity and biomass production across various taxa and ecosystems (Tilman et al. 2014, van der Plas et al. 2019, Loreau et al. 2021). Two main processes, functional complementarity and selection effects, have been proposed to explain the relationship between biodiversity and ecosystem function (BEF) (Loreau & Hector 2001, van der Plas et al. 2019, Godoy et al. 2020, Albert et al. 2022; Wang et al. 2022). Complementarity effects refer to the ability of different species within an ecosystem to perform distinct functions, leading to more efficient resource use (Loreau & Hector 2001, Poisot et al. 2013, Schneider et al. 2016). Selection effects occur when certain species are better adapted to perform certain functions. These species tend to dominate when the variation in environmental conditions favor their performance. Diversity enhances functioning and stability due to the higher probability of having species well adapted for each environmental scenario (Loreau & Hector 2001, van der Plas et al. 2019, Wang et al. 2022). The nature and strength of the complementarity and selection mechanisms could be determined by the food web in which the species are inserted (Thébault et al. 2010; Poisot et al. 2013; Wang et al. 2022; Nie et al. 2023).

The structural properties of food webs can affect ecosystem functioning along both the horizontal (i.e. within trophic levels) and vertical (i.e. between trophic levels) dimensions (Wang et al. 2017; Zhao et al. 2019). Mechanisms based on the horizontal dimension investigate how species belonging to the same trophic level allocate and compete for resources (Duffy et al. 2007, Carroll et al. 2011; Schneider et al. 2016). Communities where complementarity effects prevail, tend to exhibit high network modularity, which results from the compartmentalization of trophic interactions and spatial distributions (Rezende et al. 2009, Pinheiro et al. 2019, Borzone Mas et al. 2022), being food web modularity the multifunctional equivalent of trophic complementarity (Montoya et al. 2015). Similarly, communities where selection effects promote biomass tend to produce nested networks. In nested food webs, generalist predators consume a large portion of the resources, allowing them to dominate the communities in terms of biomass (Dattilo et al. 2014, Pinheiro et al. 2019). Then, a positive relationship between nestedness and ecosystem functioning can be expected when the dominant species significantly contribute to the ecosystem function of interest, such as standing biomass (Jiang et al. 2008, Omidipour et al. 2021). Mechanisms based on the vertical dimension examine how the different trophic levels are interrelated, through for example top-down or bottom-up controls, and evaluate the effects of interactions between predators and their prey (Bruno & Cardinale 2008, Wang & Brose 2018; Albert et al. 2022; Wu et al. 2022; Barbier & Loreau 2019, Rip & McCann 2011, Gilbert et al. 2014). The strengths of these interactions influence ecosystem functioning, where the amount of flowing matter, the efficiency of predators, their diversity and the carrying capacity of prey affect standing biomass (Gilbert et al. 2014, Nie et al. 2023). Communities with efficient consumers, and high carrying capacity of prey tend to maximize their biomass through strong interactions (H10, Figure 1), while those with inefficient predators and/or low carrying capacity of prey tend to maximize their biomass through a greater number of weak interactions (H11; Gilbert et al. 2014, Nagelkerken et al. 2020, Ullah et al. 2021).

Some mechanisms affecting ecosystem functioning can operate both at the horizontal and vertical dimensions simultaneously. For instance, niche expansion can lead to increased overlap (horizontal dimension; Chesson & Kuang 2008, Carscadden et al. 2020), resulting in more effective prey per predator ratio (vertical dimension; Burdon et al. 2019; Carscadden et al. 2020). This niche expansion and overlap can increase network connectance, which improves energy supply for predators (Thébault & Loreau 2003, Loeuille & Loreau 2005, Dunne et al. 2007), but also intensifies interspecific competition and predation effects on prey (Teng & McCann 2004, Mora et al. 2014). The relationship between connectance and biomass is variable in the literature, with positive (Thébault & Loreau 2003, Dunne et al. 2007, Lazzaro et al. 2009, Woodson et al. 2020), negative (Eschenbrenner & Thébault 2022), and independent associations reported (Danet et al. 2021). Furthermore, the coupling of different energy channels (e.g., brown and green pathways) mediated by predators helps to maintain a constant resource supply due to the asynchrony between the energy channels (Rooney et al. 2008, Rooney et al. 2012, McMeans et al. 2016, Gutgesell et al. 2022), promoting greater standing biomass (Rooney et al. 2006; Arim et al. 2010; Maureaud et al. 2019). The coupling of energy channels is determined by the connection between network compartments (Guimera & Nunes Amaral 2005, Rezende et al. 2009, Gauzens et al. 2015). A high intermodular connection by predators is expected to result in greater coupling of energy channels (Gauzens et al. 2015). Moreover, the role of intermodular connection is positively related to predator body size (McCann et al. 2005; Rooney et al. 2008; Arim et al. 2010; Pintho-Cohelo et al. 2021, Borzone Mas et al. 2022). In this sense, a greater coupling between energy channels can promote an increase in community biomass due to the constant supply of energy and/or the increase in the populations of large predators (Arim et al. 2010, Maureaud et al. 2019, Borzone Mas et al. 2022).

In summary, the BEF relationship is largely determined by the structure of food webs, which can influence biomass production through various pathways (depicted in Figure 1). However, our understanding of the mechanisms that drive biodiversity and biomass production in real-world ecosystems remains incomplete (Snelgrove et al. 2014; van der Plas et al. 2019). This is mainly due to the fact that causal pathways between these three components --diversity, food web structure and ecosystem functioning– were studied in isolation, and the challenges of acquiring relevant information from diverse and complex systems at broad scales with multiple trophic interactions (Snelgrove et al. 2014; Thompson et al. 2018, Gonzalez et al. 2020, Albert et al. 2021; van der Plas et al. 2019; Albert et al. 2022; Wu et al. 2022; Nie et al. 2023; Echenbrenner and Thébault 2023). In this work, we evaluate the relationship between richness, food web structure and standing biomass in food webs from the Paraná River floodplain in Argentina. In this sense, we will analyze how diversity constrains the structure of food webs and this affects ecosystem functioning, addressing the indirect relationships between the three ecosystem components (Figure 1).

## Materials and methods

The Paraná River belongs to La Plata River basin, which is the basin with the six largest discharges in the world and boasts around a thousand fish species (Albert & Reis et al., 2011). In its middle reach, the Paraná River has an extensive floodplain with a high diversity of microhabitats and a vast fish species richness (Marchetti et al. 2013; Scarabotti et al., 2017). The marked thermal seasonality and hydrological variability of the Parana River deeply modify the environmental heterogeneity, the riverscape connectivity, the composition of the communities and the interaction networks (Scarabotti et al. 2017; Borzone Mas et al. 2022). This combination of biodiversity, spatial heterogeneity, and temporal variability makes the Middle Paraná River an interesting model to test hypotheses about the relationship among diversity, food web structure, and ecosystem functioning (Figure 1).

Fishes were sampled from 27 water bodies grouped into four habitat types according to their flow, size and connectivity: major rivers; secondary channels, connected lakes and isolated lakes (see details in Borzone Mas et al. 2022). Sampling surveys were carried out at one of four different hydro-climatic conditions that resulted from the combination of hydrometric level and temperature seasonality in high waters cold season, high waters warm season, low waters cold season and low waters warm season. Piscivorous fish were identified, measured, and weighed. Stomachs were preserved in 10% formalin and examined in the laboratory under a binocular microscope, and prey items were identified to the species level. The biomass of each community was estimated as the total weight of captured individuals standardized by sampling effort.

### Estimation of trophic interactions and construction of food webs

Food webs were built on the basis of predator and fish prey species coexistence, predator consumption patterns, predator and prey abundances, and the energy requirements of predators. These food webs represent the sink webs of the 15 most abundant piscivorous fishes of the Middle Paraná River. Three communities with less than 3 predator species were removed from the analysis. The consumption profiles of each predator were obtained from the analysis of stomach contents and were represented as a vector with the biomass contribution of each prey to the predatoŕs diet. Co-occurrence matrices were built considering each combination of site and sampling survey (24 sites by four sampling surveys). Each fish prey species was considered as a potential resource for consumers when at least one trophic interaction with one predator was observed in any survey. The estimation of interaction strength was based on the energy required by the predator’s population, the prey preferences (consumption profile), and the functional response between prey consumption and prey abundance (see also Thébault et al. 2003, Emmerson & Rafaelli 2004; Bascompte et al. 2005), with the formula:

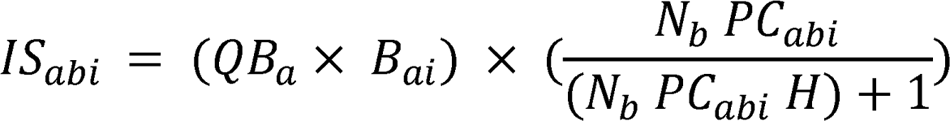

where *IS_abi_* [g / day] is the interaction strength (day biomass transfer) between the predator (*a*) and the prey (*b*) in community *i. QB_a_* [g / g. day] is the consumption rate parameter for a predator *a*, *B_ai_* [g] is the predator biomass in community i, *N_b_* [ind / ind] is the relative abundance of the prey “*b*” standardized by the abundance of all preys of predator “a’’ in the community, *PC_abi_* [g / g] is the proportion of the biomass which represents prey “*b*” on the diet of predator “*a*” in the community *i*, and *H* is the prey’s handling time (Moustahfid et al. 2010). The *QB* describes the number of times an organism consumes its own biomass in a year (Palomares & Paully 1998, Bascompte et al. 2005), here divided by 365 to infer daily interactions. The product between *QB_a_* and *B_a_* provides the biomass required by the predator population per day. In addition, because we only considered interactions among fishes, we adjusted the QB of each predator based on the proportion that fish preys represent in the diet. The second term in equation 1 represents a Holling type 2 functional response (Moustahfid et al. 2010). The consumption profile (*CPa*) is based on the proportion of biomass that each prey represents in the diet of the predator *a*. We consider that predators only feed on prey present in the community, so that *CP_ai_* is a subset of the *CP_a_* vector, with the proportions adjusted to obtain a sum equal to 1. In summary, the *CP_abi_* value is analogous to the attack rate of predator *a* on prey *b* in community *i*. The handling time (*H*) expresses the time that the predator spends on each prey and is a dimensionless parameter.

When a prey species is rare in a community or the predator shows low preference for it, the interaction strength between the two species becomes negligible, indicating a lack of significant interaction in nature. To address this scenario, we introduced a threshold value below which the interaction strength is considered zero. Since the biomass of predators and the abundance of prey vary across surveys, which significantly affects the /Sabi we calculated a specific threshold (refer to supplemental material 1). To determine this threshold, we constructed observed food webs for each of the 24 sites by pooling all recorded interactions from the samplings, and then calculated the weighted connectance (see the next section).

Under the assumption that weighted connectance remains constant within each site, we explored different values of the parameters /Sabi and H in equation 1 to find the best fit between the simulated and the observed data. To this aim, we utilized the log-likelihood function to obtain the parameter values that provided the closest match between the simulated and observed weighted connectance (Hilborn & Mangel 1997). The range of /Sabi threshold was from 0 to 1.5, at intervals of 0.01, while for handling time, the range was from 0 to 1, at intervals of 0.2. We assumed a normal distribution for the residuals between the observed and simulated weighted connectance. The combination of the /S_abi_ threshold and handling time *H* that resulted in the lowest log-likelihood was chosen for constructing the food webs for each sampling situation (refer to supplementary material 1 for methodological details and results).

### Structural properties of food webs

For each food web, we calculated species richness, weighted connectance, nestedness, interaction strength, modularity, and the strength of inter-modular connection, as follows (Banasek -Ritcher et al. 2009; Borzone Mas et al. 2022). The weighted connectance was estimated as the sum of weighted links in the network divided by the number of species (Bersier et al. 2002). Nestedness was calculated as NODF of the total matrix, a metric based on overlap and decreasing fill (Almeida-Neto et al. 2008), where values range from 0 (no nesting) to 100 (total nesting). The skewness in the interaction strength (hereafter weak interactions) examines the bias in the distribution, where positive values indicate higher proportion of weak interactions while negative values correspond to a higher proportion of strong interactions (Hair et al. 2019, see Figure 2). In this sense skewness values involve both from left to right skewed distributions from food webs with low to high diversities. Skewness was transformed by square root (Hair et al. 2019, see supplemental table 1) to control the effect of food web size. Modularity was calculated using Beckett’s method (Beckett et al. 2016), applying Newmann’s algorithm, where the modularity coefficient (*Q_w_*) was preserved. The inter-modular connector role was estimated from *c*-values (Guimera et al. 2010, Borzone Mas et al. 2022), which indicates how the species connections are distributed throughout the network, where the values range from 0 (all connections within their module) and 1 (perfect distribution among all modules in the network). Here, we used the 0.9th percentile of c-values as the maximum strength of intermodular connection, since it is less sensitive to extreme values than the maximum c-value. Since the NODF and Q_w_ values are sensitive to the size of the network, the obtained values were compared with null values from communities of the same size. In both cases, 2000 null communities were generated with fixed sums for columns and rows (quasi-swamp count) to Z-standardized values obtained based on null model expectations. See Supplementary table 1 for the functions and packages used to compute the structural variables.

**Figure 2:**
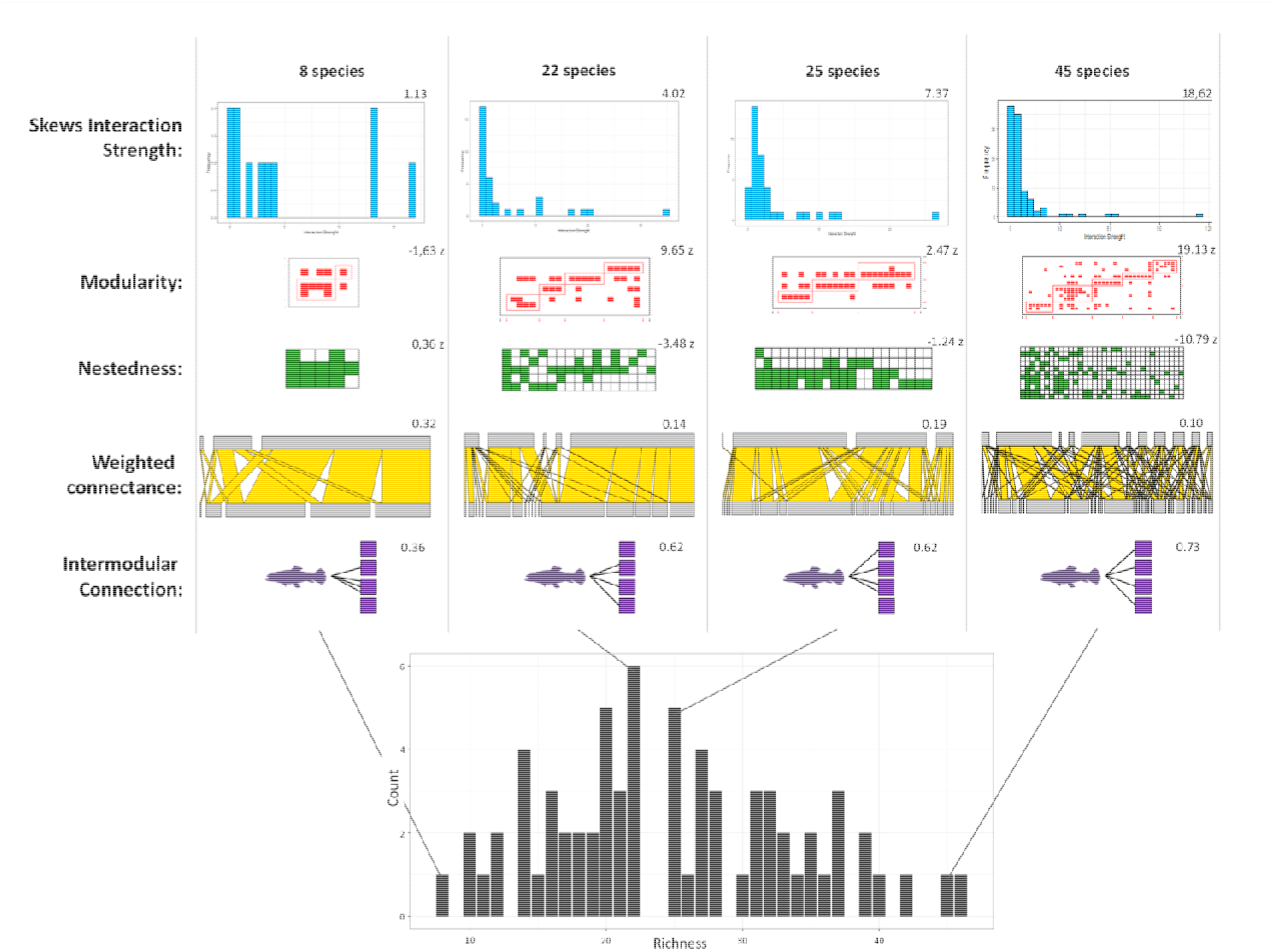
Changes in the structure of the food web based on richness. Each community is represented by different aspects: weighted connectance (yellow), modularity (red), nestedness (green), frequency of interaction strength (blue), and intermodular connection (violet). A) Frequency distribution of fish community richness in the Paraná River. Four examples of food webs are presented, illustrating a gradient of richness: 8 species, 22 species, 25 species, and 45 species. As richness increases, weighted connectance and nestedness decrease while, modularity, the frequency of weak interactions, and modular connection by predators increase with the number of species. The representation of the intermodular connection is schematic, where the distribution of the connections between the modules corresponds to the c-values in each food web, but the number of modules and connections is simplified.

To test our twelve hypotheses, we analyzed the relationships among species richness, food-web structure and community biomass with a path analysis (Shipley 2016; Borthagaray et al. 2020). Starting with the causal structure of Fig. 1, non-significant relationships were deleted and whole model performance was evaluated until we obtained a causal structure with significant paths supported by data. Discrepancies between observed and expected covariance matrices were assessed with ^2^ tests. To estimate the fit of the whole model we used the root mean square error approximation (RMSEA)(Shipley 2016). RMSEA values close to zero indicate good model fit. The Lavaan package of R was used in these analyses (Rosse 2012). All variables were centered and standardized to eliminate the differences in magnitude.

## Results

A total of 21,232 individuals belonging to 147 species were recorded. The 15 piscivorous species accounted for 1,249 trophic links. Sixty three food webs were built ranging from 5 to 38 species. The thresholds of /Sabi and handling time (*H*) differed among surveys ranging from 0.06 to 0.39 (Supplemental material 1). Values of richness, food web structure and biomass for each community can be found in supplementary table 2. After removing the causality links in figure 1 among nestedness, weak interactions and inter-modular connection with biomass, we obtained a causal model supported by data (Fig. 3; X^2^= 2.60; p-value = 0.62, N = 63; RMSEA= 0.024, 90 % upper = 0.15; 90% lower = 0.000).

**Figure 3:**
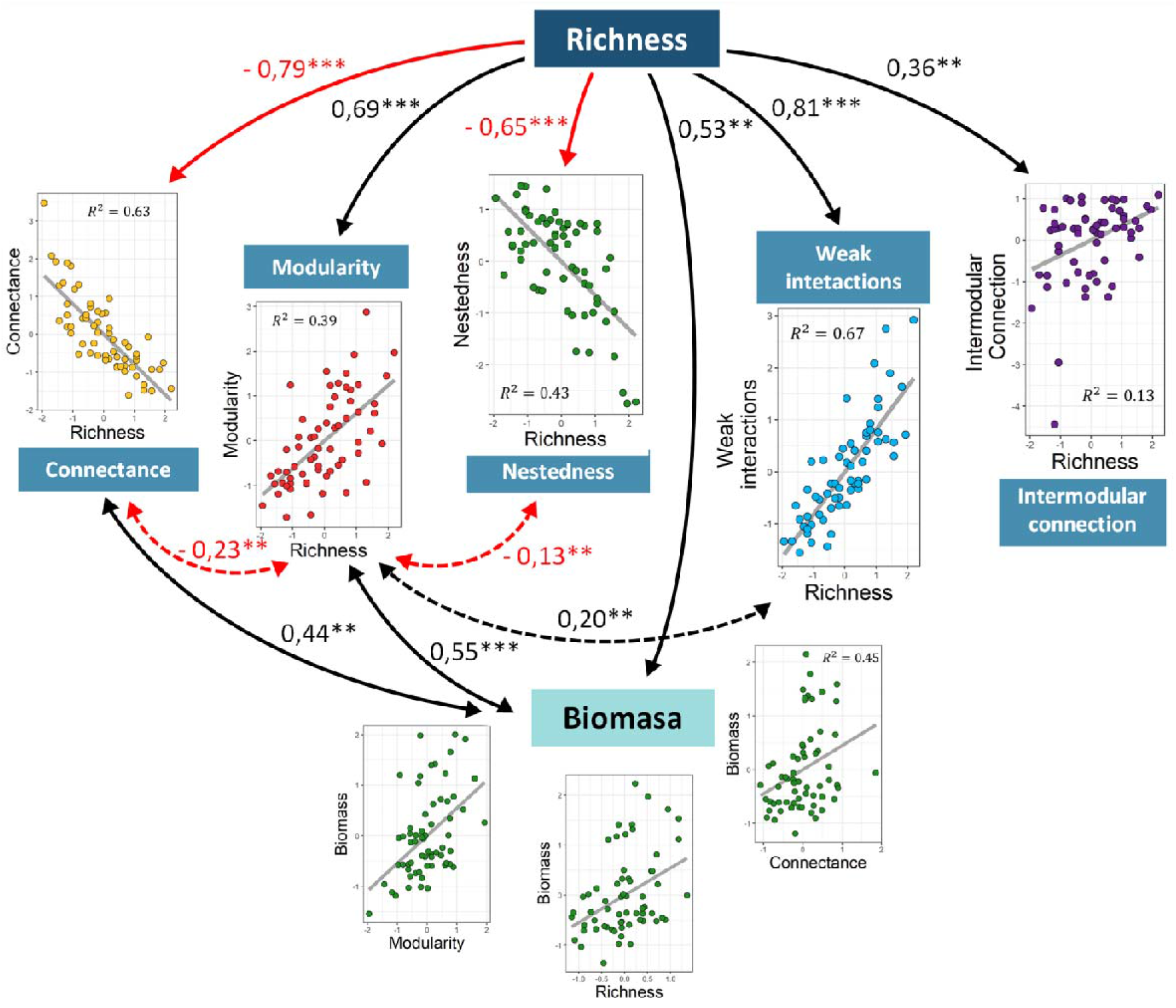
Significant relationships between richness, food webs structures and biomass. The bivariate plots were created with partial residuals after removing the effects of all the other paths. The black arrows indicate positive relationships, while the red ones indicate negative associations. Directional arrows show causal relationships, while bidirectional arrows represent associations without causality (covariance). The asterisks on the lines express the significance of the relationship, where * < 0.05; ** < 0.01 and *** < 0.001 and the coefficients are the slope of the regression. Values of R2 within the plots indicate the amount of variation in the variable that is explained by the path model.

All the variables of food web structure showed a significant relationship with richness, which generally explained a large proportion of the variation in food web metrics (Fig. 3). The number of weak interactions, modularity and inter-modular connection ability increased with species richness, whereas connectance and nestedness decreased. In turn, biomass was positively related to richness, modularity and connectance. Finally, richness showed two indirect pathways towards biomass, a negative one mediated by connectance (total effect: −0.34) and a positive one mediated by modularity (total effect: 0.37). Nestedness and modularity, as well as modularity and connectance were negatively correlated (Fig. 3).

## Discussion

This study advanced in the clarification of the relationship between three key components of ecosystems: their diversity, their food web structure, and their functioning. Firstly, we report that food web properties consistently change along a species richness gradient as predicted by theory. This includes decreasing connectance, nestedness and trophic interaction strength, while increasing modularity and the role of some species connecting different trophic modules with the increase in species richness (McCann 2011; Rooney & McCann 2012; Maureaud et al. 2020). The observation of the direct effects of community richness in each one of the food web metrics, while controlling the relationships between food web metrics, represents a significant advance in the empirical support of food web theory. The causal structure supported here is one of the several ones congruent with actual theory (e.g. Danet et al. 2021; Eschenbrenner & Thébault 2023). Secondly, when we statistically controlled the indirect effects of food web structure on biomass by path analysis, we observed a direct effect of species richness on the standing biomass. Thirdly, we also found large indirect effects of species richness on the community biomass mediated by its effects on food web modularity and connectance. Notably, this involves a positive effect due to the increase in modularity but also a negative effect mediated by the decay in connectance with community richness. All these results support the view that the effect of biodiversity on community biomass and food web structure are intertwined (Poisot et al. 2013, Wang and Brose 2018, Daam et al. 2019, Wu et al. 2022, Nie et al. 2023).

A better understanding of the effects of structural properties of ecological networks on biodiversity, stability, and functioning has been a significant advancement in ecology of the last decades (Stouffer et al. 2010; Valdovinos et al. 2010; Landi et al. 2018; Damm et al. 2019; Goyal et al. 2022; Ratzke et al. 2020). In particular, an effect of community richness on food web connectance, nestedness, modularity, intermodule connectivity and interaction strength is consistently supported both in theoretical and recent empirical studies. However, less attention was devoted to the interrelationship of these metrics along diversity gradients. Multiple causal structures are compatible with the diversity-complexity relationship in food webs. Here, we identify a causal structure well-supported by data, in which species richness has a strong direct effect on each one of the food web metrics considered, in addition to a complex network of causal relationships among food web metrics These relationships suggests that there are multiple mechanisms by which community assembly responds to richness gradients, putatively allowing communities to be diverse and stable.

Our findings demonstrate that an increase in species richness promotes higher modularity and lower connectance and nestedness in ecological networks. The persistence of these structural features in natural interaction networks is considered to be associated with the topological stabilization, avoiding the propagation of disturbances and reducing the variability of whole systems (Stouffer and Bascompte 2010; Welti & Joern 2015, Gillarranz et al. 2015, Ceron et al. 2019, Eschenbrenner & Thébault 2023, but see Kondoh et al. 2010, Thébault and Fontaine 2010). Furthermore, in heterogeneous environments, different species might be forced to use different microhabitats and consume different preys promoting an aggregation of interactions in different modules (e.g. Borzone Mas et al., 2022). A strong heterogeneity in habitat types or feeding resources can additionally reduce the performance of generalist species in favor of specialist species because they are unable to exploit resources along such a wide range of resources (Pinheiro et al., 2019). These mechanisms determine a modular aggregation of interactions in heterogeneous environments (Pinheiro et al., 2019; Borzone Mas et al., 2022). The Paraná River floodplain harbors 330 fish species with broad body size distribution and diverse ecomorphological characteristics (e.g. spanning 5 orders of magnitude in body weight), and life history strategies (Scarabotti et al. 2011, 2017, Borzone Mas et al. 2019, Abrial et al. 2019). In this system, functional diversity in fish prey imposes constraints on trophic interactions, such as gape size limitation, energetic return limitation, barrier traits, and forbidden interactions (Arim et al. 2010, Ortiz & Arim 2016, Izquierdo-Palma et al. 2021). The required match in predator and prey traits promotes aggregation of interactions among species interacting in particular environments, of consumers guilds in some resources, and of prey species of predator guilds, promoting a direct association between modularity and species richness (Sherwood et al. 2002, Pinheiro et al. 2019). In contrast, studies in communities with narrow body size distribution, low ecomorphological diversity, and homogeneous life history strategies have reported positive associations between species richness and nestedness/connectance, due to the lack of constraints on trophic interactions (Kondoh et al. 2010, Thébault et al. 2010, Pinheiro et al. 2019, Ho et al. 2022).

It should be noted that the complexity-stability literature, formerly relating richness, connectance and interaction strength (May 1972), devoted central attention to the richness-connectance relationship with less attention to the richness-interaction strength association (Yodzis 1981, McCann et al. 1997, Paine 1992, Berlow et al. 2009; Gross et al. 2009). The importance of weak interactions in ecological networks for community stability, such as damping oscillations in abundance and providing alternative resources in changing environments, is well accepted (Quince et al. 2005, Berlow et al. 2009; Loueille 2010; Rooney & McCann 2012). However, the prediction that the biases toward weak interactions increase with richness (Berlow et al. 2009; Gross et al. 2009) has been rarely considered in empirical studies. Our results confirm the dominance of weak interactions in the food webs of the Paraná River, but also highlight that the biases toward weak interactions is tightly associated with community richness. In addition, the presence of inter-modular connections by species feeding in alternative compartments of food webs, which is theoretically supported as a stabilizing property and has been progressively reported in different ecosystems (McCann 2011; McMeans et al. 2016, Borzone Mas et al. 2022), was found to significantly increase with diversity in our study. This relationship could arise as a by product of the increase in modularity, which makes it more likely for predators to capture prey from different modules. However, our path analysis (Fig. 3) did not show a direct association between modularity and inter-modular connectivity; indicating that the increase in inter-modular connection is a genuine effect of the increase in species richness, more than just a byproduct of the rise in modularity. As in other systems, the role of modular connectors in the Paraná River food web is determined by predator traits such as body size and ecomorphology, (Rezende et al. 2009; Pinto-Coelho et al. 2021; Borzone Mas et al. 2022). Probably the change in the representation of these traits in diverse communities is involved in the direct effect of richness on inter modular connectivity.

Food web structure and ecosystem functioning dominated by cascade effects, focused on diversity and biomass of consumers or primary productivity, but seldom considering other metrics (Carpenter et al. 1987; Polis & Strong 1995; Hooper et al. 2005). We found that standing biomass is positively correlated with richness, modularity, and connectance (Figure 3). Modularity might promote higher trophic complementarity and niche packing, resulting in more efficient distribution of resources within and between modules, which in turn promotes higher biomass production (Montoya et al. 2015, Schneider et al. 2016, Wang & Brose 2017). An increase in interactions can lead to two alternative mechanisms promoting standing biomass. On one hand, a high number of interactions maintains the supply of resources to predators (Gilbert et al. 2015). On the other hand, a higher degree of overlap can increase negative interspecific interactions between predators, leading to self-regulation within consumer populations, which can have positive impacts on prey populations and food web stability (Teng & McCann 2004, Mora et al. 2014, Woodson et al. 2020, Zhao et al. 2019). Previous studies did not find a significant association between connectance and biomass in stream fish (Danet et al. 2021). However, while this study proposed a similar conceptual framework to the present one, they did not analyze the role of modularity. In our framework, the relationship between connectance and biomass was evidenced when modularity was incorporated into the model (see supplementary material 2). This suggests that considering the number of interactions without taking into account how they are distributed (e.g., in modules) may provide a biased idea about the effect of connectance on ecosystem functioning.

In this study, we add empirical evidence supporting that the structure of the food web has to be considered for fully understanding the mechanisms underlying the BEF relationship (e.g. Wang and Brose 2018; Nie et al. 2023; Albert et al. 2022; Eschenbrenner and Tébault 2023; Wu et al. 2023). Indeed, the considerable variation in the diversity-productivity relationship that has been reported may be related with differences in the food web scenarios in which the BEF was analyzed (see also Wu et al. 2022; Nie et al. 2023).

The causal hypotheses supported by our data (Figure 1 and 3) suggest that species complementarity – linked to modularity– and the number and efficiency of energy channels– linked to connectance–play a crucial role in shaping the relationship between diversity and standing biomass in the Paraná River. These findings align with other research in aquatic systems, which has also observed that complementarity in the horizontal dimension (Gilarranz et al. 2015, Woods et al. 2020, Danet et al. 2021, Moi et al. 2021) and the control of prey populations by predators or cascade effects in the vertical dimension (Fung et al. 2015; Danet et al. 2021, Yamamuro et al. 2019) can promote community biomass (see also Maureaud et al. 2019, Fu et al. 2021, Feng et al. 2022). As a consequence, changes in environmental conditions or habitat degradation that disrupt food web connectance or modularity (or their relative values) can alter the mechanisms that shape the association between community diversity and standing biomass. For example, landscape homogenization, reduction in the range of body sizes, or the invasion of a generalist predator can disrupt the relationship between species richness and modularity, resulting in decreased biomass production (Rezende et al. 2009, Montoya et al. 2015, Kovalenko 2019). In such cases, when modularity conditions are lost, an increase in community diversity amplifies the negative indirect effects on biomass due to the adverse impact of diversity on connectance. This implies that human activities that degrade the structural properties of food webs can disrupt the mechanisms that shape the BEF relationship, and the ecosystem services provided to humans (Moi et al. 2023). Consequently, sustaining diverse and productive communities requires the presence of suitable conditions for a modular organization of food webs.

A pressing priority in ecology is to develop a mechanistic understanding of how global change impacts biodiversity and ecosystem functioning (Akesson et al. 2021). Our study provides empirical evidence for the connection between diversity and food web structure, as well as, about the effect of community richness on standing biomass and the indirect effects mediated by specific features of food webs–in this case connectance and modularity (Worm & Duffy 2003; Arim et al. 2010, Poisot et al. 2013, Wang and Brose 2018, Albert et al. 2022; Wu et al. 2022, Nie et al. 2023). Therefore, solely examining the direct association between species richness and biomass without considering changes in food web structure may provide a limited understanding of the real connection between diversity and ecosystem functioning (Poisot et al. 2013, Wang and Brose 2018, Daam et al. 2019, Wu et al. 2022, Nie et al. 2023). Our findings support the notion that mechanisms operating simultaneously in both the horizontal and vertical dimensions shape ecosystem functioning (Wang & Brose 2018; Zhao et al. 2019; Wu et al. 2022). Importantly, our results also suggest that human activities that promote environmental homogenization, changes in species traits, and species pool dynamics through biological invasions and extinctions can undermine the key mechanisms that connect diversity, food web structure, and ecosystem functioning, leading to potential destabilization of ecosystem functioning. In this way, conservation efforts should consider the impacts of human activities on food web structure if the ecosystem functioning is to be preserved.

## Box text figure 1)

Richness ∼ Biomass (positive): Three main mechanisms shape the relationship between biodiversity and standing biomass, these are complementarity effects, selection effects and the sampling effect, although many authors consider the last two as a single phenomenon (Loreau 2010; Tilman et al. 2014, van der Plas 2019).

Richness ∼ connectance (negative): This expectation is supported by four mechanisms: i.-trophic network stability, the overconnected food webs tend to be unstable (May 1972; Stouffer and Bascompte 2011) ii.-sampling effects, the fraction of detected interactions decrease with the size of the web, (Warren 1994), iii.-spatial mismatch, in larger food webs increase the fraction of predators and prey that do not inhabit the same microhabitats; and iv.-biological constraints, the trait matching promote forbidden interactions (May 1972; Warren, 1994; Thebault et al., 2010). Species interacts with a fixed average number of species not increasing with the community richness (Cohen 1990) or increasing at lower rates than the total number of possible interactions (Martinez 1992) determining a reduction in connectance with the increase in species richness.

Richness ∼ Modularity (positive): Several mechanisms may promote an increase in modularity with community richness: i) modularity reduce connectance and the average interaction strength, representing a feasible architecture at large diversity levels (May 1974: Stouffer and Bascompte 2011); ii) spatial compartmentalization, community diversity is usually associated with the heterogeneity of microhabitat, spatially aggregating species and interactions (Pillai et al 2012; Canavero et al. 2014; Borzone-Mass et al. 2022); iii) resource heterogeneity in diverse communities determine the concentration of interactions in different kinds of resources or predatory strategies Rezende et al. 2009; Gillarranz et al. 2015; Pinheiro et al. 2019); iv) discontinuities in predator-prey trait distributions determine the cluster of interactions where traits matches are more likely; these discontinuities could both increase or decrease with diversity (Borthagaray et al. 2012; Sherwood et al. 20011, 2012) determining an increase or a decrease in modularity respectively.

Richness Nestedness (Negative): Nestedness and modularity are, at some level, two sides of the same coin (Fortuna et al. 2010). Consequently, all the above mentioned mechanisms that promotes an increase in modularity with diversity, may also promote a decrease in nestedness. In addition, in systems with a high resource diversity might foster a decrease in nestedness due to a reduction in the fraction of total prey number consumed by each predator (Pinheiro et al. 2019; Cerón et al. 2019). However, a gradient in consumers traits—e.g. body size—determines a gradient of optimal resources to be consumed—e.g. zooplankton, macroinvertebrate, fishes; Ritchie 2010; Sherwood et al. 2011, 2012; Arim et al. 2011). In communities with high diversity, prey are progressively incorporated by predators of larger size determining a nested arrangement of trophic interactions (Arim et al. 2010), but low prey diversity stack consumers in alternative prey categories promoting modularity (Sherwood et al. 2011, 2012).

Richness ∼ weak Interactions (Positive): As prey diversity increases, interactions strength biases toward weak that represent a feasible pattern due to its effect in stabilizing species coexistence and ensuring resource availability (Yodiz 1981; McCann et al. 1997, Berlow et al. 2009). In addition, strong functional responses towards a specific type of prey may be inefficient in diverse communities, leading predators to broaden their niche breadth and reduce the strength of interaction with each prey (De Long & Coblentz, 2021).

Richness ∼ Intermodular connection (Positive): Species richness is associated with the number of energy channels and resources (Arim et al. 2010). Consequently, high prey richness implies greater compartmentalization in interactions, increasing the potential intermodular connection capacity of predators (Arim et al. 2010, Rooney et al. 2012).

Connectance ∼Biomass (Positive): The increase in connectance is related to a higher level of supply and efficiency of the network, and networks with generalist predators manage to incorporate a greater number of resources and increase the self-regulation between of predators. (Teng & McCann 2004; Dunne et al. 2007, Loreau et al. 2010).

Modularity ∼ Biomass (Positive): Modularity emerges as a multifunctional equivalent of the complementarity effect. Module-distributed interactions promote niche packing, decreasing interspecific competition and apparent competition between predators and prey from different compartments respectively. More modular networks allow greater efficiency in the use and distribution of resources in energy channels, promoting community biomass (Montoya et al. 2015; Gillarranz et al. 2015; Moi et al. 2015).

Nestedness ∼ Biomass (Positive): Nested food webs provide an advantage in resource use for generalist species. If these species contribute to the ecosystem function analyzed (for example biomass accumulation), a positive association between nestedness and ecosystem functioning is expected. In turn, it can emerge from communities where selection effects operate (Fu et al. 2021, Feng et al. 2022).

Strength interactions ∼ biomass (Negative): Weak interactions provides alternative energy path in variable environment, allowing species to preserve high biomass levels (Berlow et al. 2009). In addition, inefficient predators with weak interactions toward their prey may foster coexistence but allowing prey guilds to be close to their carrying capacity, promoting community biomass (Gilber et al. 2014, Wang et al. 2019) Strength’s interactions ∼ Biomass (positive): In situations where predator efficiency is high and prey carrying capacity is high, strong interactions towards large predators promote community biomass. Under these conditions, a higher flux towards predators with lower metabolic expenditure allows for an increase in biomass, however, this maximization of standing biomass through strong interactions often undermines food web stability (Gilbert et al. 2014, Nie et al. 2023).

Intermodular connection ∼ Biomass (Positive): The coupling between of food web modules by consumers enhance these consumer biomasses by means of ensuring the availability of alternative resources and expanding the energetic support for predators (Arim et al. 2010, Rooney et al. 2012).

