## Supplemental table 1 for "Food web structure mediate positive and negative effects of diversity on ecosystem functioning in a large floodplain river"

| variable | Function | Package |
| --- | --- | --- |
| wieghted connectance | networklevel(fwmat, weighted = TRUE, index = c("weighted connectance")) | bipartite |
| modularity | metaComputeModules(fwmat, N = 5)@likelihood | bipartite |
| NODF | oecosimu(fwmat,nestednodf, "r2dtable") | vegan |
| c-value | czvalues(mod, level = "higher") | bipartite |
| skewness interactions strengths | skew(linksreal$value)/sqrt(6/nrow(linksreal)) | phych |

**Supplemental table 1:** Description of the functions used in rstudio and the packages to calculate the structural variables of food webs. Among the functions it can be observed fwmat = trophic matrix for each community, mod = moduleWeb object created from the metaComputemodules function and linksreal = list with the strength of each trophic interaction of the local trophic web. Furthermore, r2dtable is a randomization algorithm which maintains the marginal sums of rows and columns.
