## Supplemental material 1 for "Food web structure mediate positive and negative effects of diversity on ecosystem functioning in a large floodplain river"

When a prey is rare in a community or the predator has low preference for it, the interaction strength between the two species is negligible, indicating no significant interaction in nature. To account for this situation, we introduced a threshold value below which the interaction strength is considered zero. To obtain this threshold, we built food webs with trophic interactions collected in the field. In this sense, 24 real food webs were built grouping the interactions observed in each site along all sampling surveys. To adjust the threshold value of interaction strength SI_bai_ and handling time (H), the likelihood method was used (Hilborn & Mangel 1997). In this sense, the weighted connectance of the real food webs was calculated and related to the weighted connectance of simulated food webs built from equation 1 (see reference in the main text).

$${Si}_{bai}={QB}_{b}\times B_{bi} \times\frac{N_{a} \times{cp}_{api}}{\left( N_{a} \times{cp}_{api} \times H \right)+1}$$

The weighted connectance considers the network connectance as a function of the number of species, allowing the elimination of the effect of richness in the comparison between real and simulated food webs (Bersier et al. 2002, Banasek-Ritcher et al. 2009). Then, we calculated the numerical values of Si_bai_ and H that best fitted the weighted connectance between simulated food webs and real food webs. Si_bai_ was analyzed between 0 and 2, at intervals of 0.01, while H was evaluated between 0 to 1, at intervals of 0.2. In each combination of parameters, we observed the log-likelihood values of the adjustment of the connectance between real food webs and simulated food webs, and selected the model with the lowest log-likelihood value (Hobbs & Hilbron 2006). We assumed a normal distribution for the residuals between the observed and simulated weighted connectance. This procedure was carried out for each sampling survey; therefore, the threshold value of the Si_bai_ and the parameter H differ in each hydroclimatic condition.

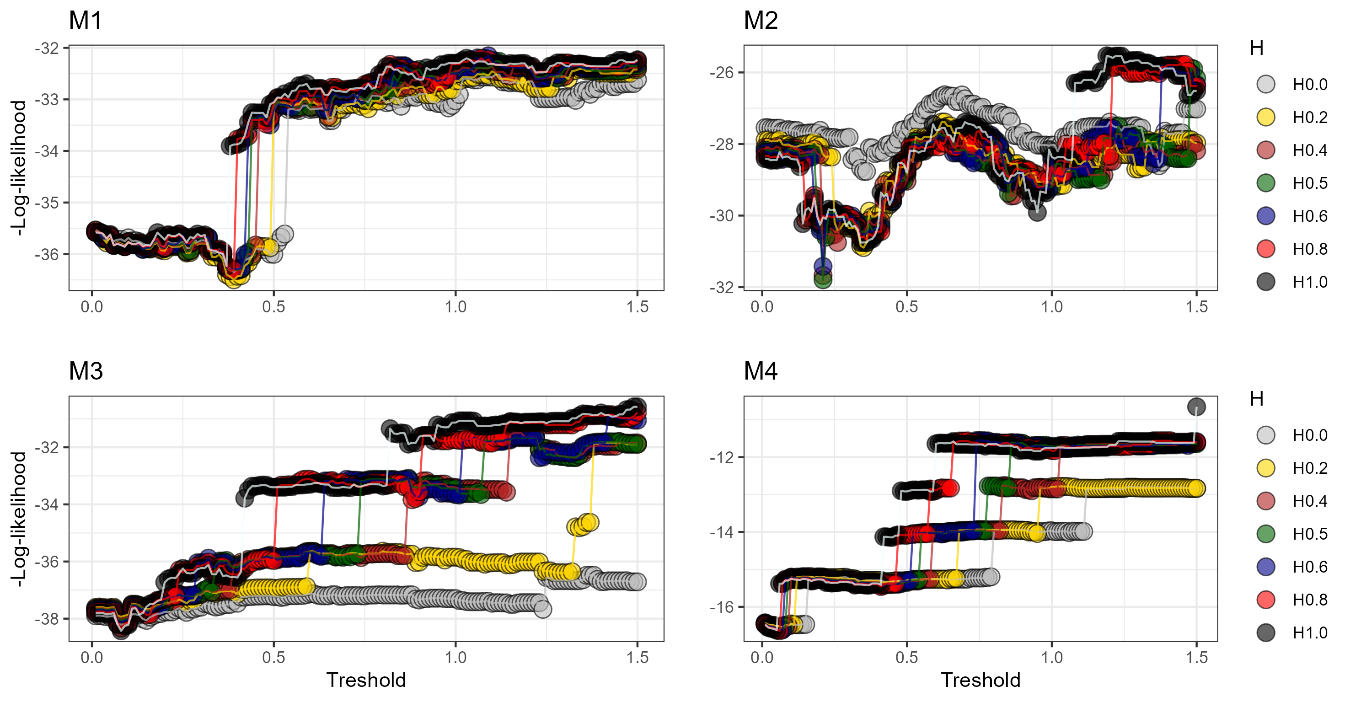

Each graph corresponds to a different sample. The x-axis shows the threshold value of the interaction force, while the Y-axis shows the – log-likelihood- value. The more negative the log-likelihood values are, the better fit is observed between the results of the model and the values shown

Within each graph, each color of lines and points represents the value of the parameter H (handling time). The parameters selected for the model in each sampling can be seen below

| Sample | Interaction Strength | Handling Time | -log-likelihood |
| --- | --- | --- | --- |
| M1 | 0.39 | 0.2 | -36.5 |
| M2 | 0.21 | 0.5 | -31.8 |
| M3 | 0.08 | 0.4 | -38.42 |
| M4 | 0.06 | 0.8 | -16,14 |
