## Supplemental material 2 for "Food web structure mediate positive and negative effects of diversity on ecosystem functioning in a large floodplain river"

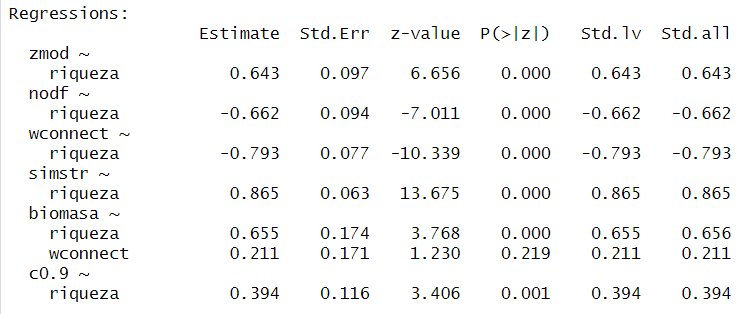

Pathway analysis coefficients similar to figure 3 after removing the causal relationship between modularity and biomass. As observed in the coefficient highlighted in red, the relationship between connectance and biomass is not significant (p-value = 0.21).
